# Searching for intra-locus sexual conflicts in the three-spined stickleback (*Gasterosteus aculeatus*) genome

**DOI:** 10.1101/2022.07.04.498749

**Authors:** Florent Sylvestre, Claire Mérot, Eric Normandeau, Louis Bernatchez

## Abstract

Differences between sexes in trait fitness optima can generate intra-locus sexual conflicts that have the potential to maintain genetic diversity through balancing selection. However, these differences are unlikely to be associated with string selective coefficients and as a result are challenging to detect. Additionally, recent studies have highlighted that duplication on sexual chromosomes can create artefactual signals of inter-sex differentiation and increased genetic diversity. Thus, testing the relationship between intra-locus sexual conflicts and balancing selection requires a high-quality reference genome, stringent filtering of potentially duplicated regions, and dedicated methods to detect loci with low levels of inter-sex differentiation. In this study, we investigate intra-locus sexual conflicts in the three-spined stickleback using whole genome sequencing (mean coverage = 12X) of 50 females and 49 males from an anadromous population in the St. Lawrence River, Québec, Canada. After stringent filtering of duplications from the sex chromosomes, we compared three methods to detect intra-locus sexual conflicts. This allowed us to detect various levels of inter-sex differentiation, from stronger single locus effects to small cumulative or multivariate signals. Overall, we found only four genomic regions associated with confidence to intra-locus sexual conflict that displayed associations with long-term balancing selection maintaining genetic diversity. Altogether, this suggests that most intra-locus sexual conflicts do not drive long-term balancing selection and are most likely transient. However, they might still play a role in maintaining genetic diversity over shorter time scales by locally reducing the effects of purifying selection and the rates of genetic diversity erosion.

## Introduction

In species with sexual reproduction, sexes often have different fitness optima for shared traits, resulting in sex-specific selection pressures (Lande 1980; Rowe et al. 2018). If males and females share the same genetic basis for such a trait, natural selection can favour different alleles in each sex, leading to intralocus sexual conflict (Van Doorn 2009). Because sexual reproduction randomly shuffles alleles between sexes at each generation, such conflicts have the potential to generate balancing selection, maintaining male and female-beneficial alleles in the population, depending on the selective coefficients in each sex (Connallon and Clark 2014; Lonn et al. 2017; Zajitschek and Connallon 2018). Intra-locus sexual conflicts have been identified at the phenotype level in a variety of taxa and traits (Merilä et al. 1997; Foerster et al. 2007; Cox and Calsbeek 2009; Harano et al. 2010). As such, they have the potential to play an important role in maintaining genetic diversity in natural populations.

Direct detection of intra-locus sexual conflict implies measuring fitness in male and female samples and identifying genotypes with opposite fitness effects between sexes (Innocenti and Morrow 2010; Ruzicka et al. 2019). Yet, accurate fitness measurements are at best difficult to obtain in natural populations. Another solution is to search for differences in allele frequency between males and females caused by sex-specific mortality (Rowe et al. 2018). Several recent studies have used this approach to identify intra-locus sexual conflicts in large-scale genomic datasets (Cheng and Kirkpatrick 2016; Lucotte et al. 2016; Dutoit et al. 2018; Wright et al. 2018; Wright et al. 2019). More recently, Kasimatis et al. (2021) failed to find signal of such conflict despite using dataset with much more statistical power (N = 409,406 human genomes. Confidently identifying signal of intra-locus sexual conflict thus remains a challenging task.

Since sexual reproduction erases genetic differences between males and females except on sex chromosomes, differentiation caused by sex-specific mortality is expected to be weak. Moreover, substantial mortality in each sex is required to reach a significant detection level (Kasimatis et al. 2019; 2021). It is unlikely that most intra-locus sexual conflicts over survival are under strong selective pressure as this would represent too high a mortality cost for populations to persist. Given the low magnitude and detection power involved in intra-locus sexual conflicts, separating signal from noise is a complex task, especially in large and complex whole genome sequencing datasets in which multiple testing allows detection of only the strongest signals. One way to overcome this challenge may be to search for cumulative signals of differentiation over several loci, as opposed to looking for significance at a single locus. This approach improves the detection power of intra-locus sexual conflict in natural populations and makes detecting smaller selective effects possible (Ruzicka et al. 2020).

Another factor to consider is that sequence similarity between autosomes and sex chromosomes can generate confounding signals of intra-locus sexual conflict (Kasimatis et al. 2019; Bissegger et al. 2020; Lin et al. 2022). If a locus is duplicated in a sex-specific genomic region, it can accumulate sex-specific polymorphism (Bissegger et al. 2020; Mank et al. 2020; Lin et al. 2022). If the region remains similar to its original autosomal copies, it is possible that sequencing reads from the sex-specific region align to the autosomal copies, creating a spurious signal of differentiation between sexes that can be mistaken for an intra-locus sexual conflict. Moreover, duplications of loci on sexual chromosomes are also expected to be part of the process of resolving intra-locus sexual conflict (Rice 1984). This could lead to signals of intralocus sexual conflict that may in fact be caused by resolved sexual conflict (Mank et al. 2020; Lin et al. 2022). The use of a high-quality reference genome with identified sex chromosomes and stringent filtering for potential duplicates is thus essential for the reliable detection of intra-locus sexual conflict.

It is still unclear what proportion of intra-locus sexual conflicts play a role in maintaining genetic diversity in natural populations. Their contribution to genetic diversity depends on the selective coefficients involved (Zajitschek and Connallon 2018), but also on the ultimate resolution of such conflicts. Several studies suggest that intra-locus sexual conflicts might be easily resolved by the evolution of a sex-specific genetic basis for a given trait or differential gene expression (Wright et al. 2018; van der Bijl and Mank 2021). In such a scenario, intra-locus sexual conflict would not be associated with longterm balancing selection but rather could affect genetic diversity by locally reducing the impact on background selection around the selected loci until the resolution or the end of the sexual conflict (Ruzicka et al. 2020). To better understand the respective roles of these alternative mechanisms, we need to rigorously test the association between long-term balancing selection and intra-locus sexual conflict.

Here, we aim to investigate the presence of intra-locus sexual conflicts in a natural population of three-spined stickleback (*Gasterosteus aculeatus*). This model species presents several advantages for the study of intra-locus sexual conflict: 1) sticklebacks exhibit pronounced sexual dimorphism for a variety of traits (e.g.: behaviour, colouration, reproductive strategy; Whoriskey et al. 1986; Chellappa and Huntingford 1989; Barker and Milinski 1993); 2) they produce large juvenile cohorts (Craig and FitzGerald 1982; Poulin and Fitzgerald 1989), thus having the potential for a high juvenile mortality load; 3) a very high-quality reference genome (Nath et al. 2021) with a recently sequenced Y chromosome (Peichel et al. 2020) is available, which mitigates the issue of duplicated genes on sex chromosomes.

We performed whole-genome sequencing of 49 male and 50 female three-spined stickleback to search for signatures of intra-locus sexual conflicts. We first used sex-specific variation in coverage and genetic variability to filter potentially duplicated variants on sex chromosomes. Then, we searched for signals of intra-locus sexual conflicts throughout the genome by combining single- and multi-locus approaches, aiming to detect loci with weak polygenic effects. Finally, we tested for the association between potential intra-locus sexual conflicts and genetic diversity and potential balancing selection.

## Results

### I Investigating intra-locus sexual conflicts requires adequate filtering

Artefactual signatures mimicking intra-locus sexual conflicts may be created by both population factors (sampling, structure, inbreeding) and genomic factors (sex chromosomes, rearrangements, duplications). To minimize the impact of such potential artefacts, we studied a single population and applied stringent filtering. First, to avoid confounding effects due to population factors, we sampled and sequenced a total of 100 anadromous three-spined stickleback (50 males and 50 females) from a single panmictic population from the St. Lawrence R. Estuary (McCairns and Bernatchez 2009). After stringent filtering, we genotyped 1,755,558 autosomal biallelic SNPs (see methods and table S1 for filtering details). One male shown elevated missingness compared to other and was discarded. No population structure nor sex-specific structure was apparent on a PCA, confirming that our samples belong to a single panmictic population (Fig. 1A, Fig S1). Investigating pairwise kin coefficients between samples, we confirmed that relatedness among individuals was equivalent between males and females (Fig 1B) and should therefore not influence inter-sexual differentiation. Second, to avoid artefacts due to genomic factors, we excluded SNPs located in large structural variants from our dataset. Local PCAs generated by sliding windows identified three regions on chrIX (4.75Mb), chrXVI (0.95Mb) and chrXXI (0.90Mb) that displayed characteristics consistent with polymorphic inversions and potential duplication (Fig S2, S3 and S4, Supplementary discussion). However, those structural variants showed neither sex bias nor inter-sex differentiation (Supplementary discussion) and are thus unlikely candidates for intra-locus sexual conflicts.

**Figure 1:**
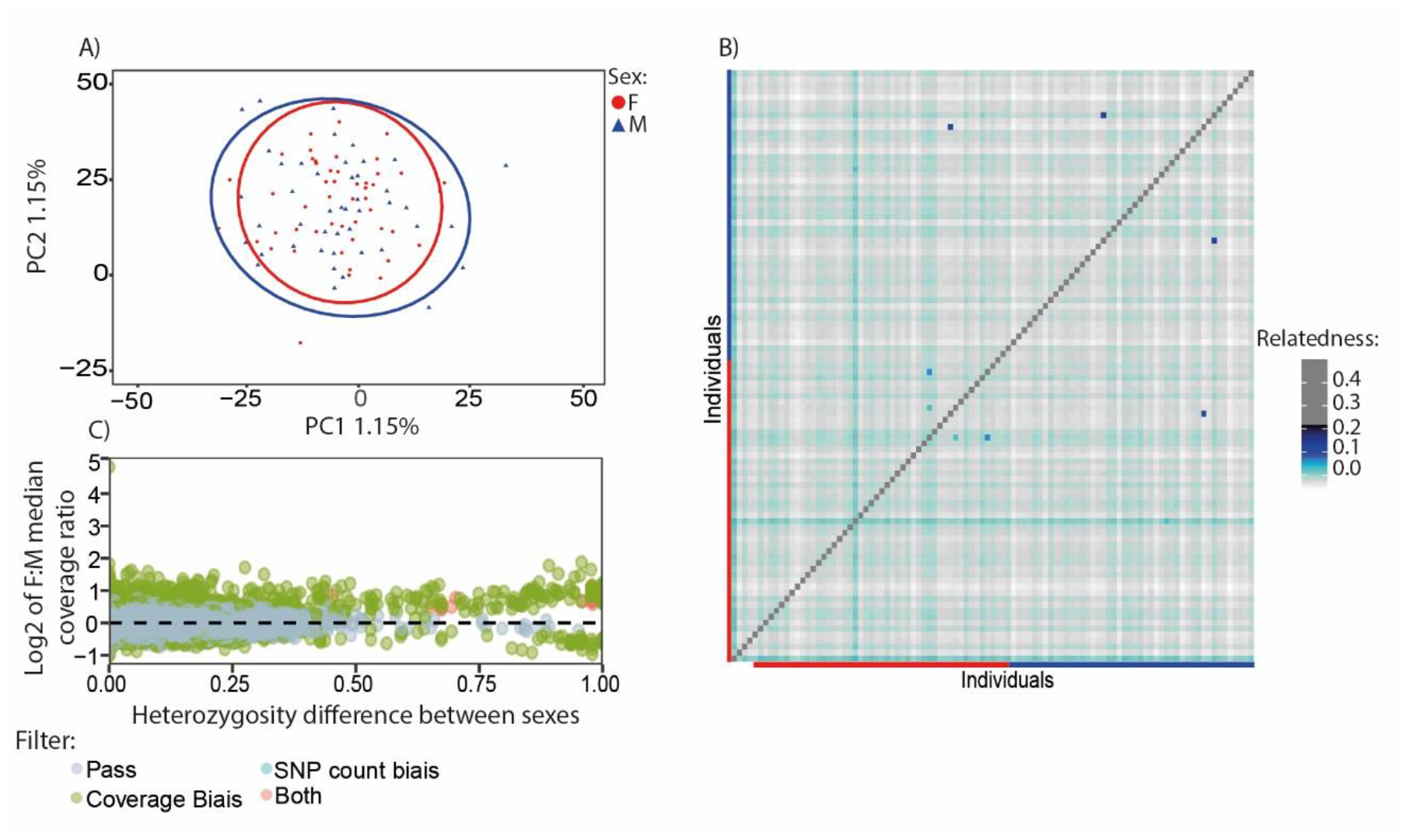
Global patterns of inter-sex differentiation and duplications on sexual chromosomes. A) First two axes of a Principal Component Analysis (PCA). Two outlier individuals with high relatedness are not shown. Ellipses represent the 95% confidence interval of point distribution. Red (circle and ellipse) represent females, blues (triangle and ellipse) represent males. B) Heatmap of pairwise relatedness as estimated by the KING implementation in vcftools. C) Variation of female/male coverage ratio as a function of absolute heterozygosity difference between sexes for autosomal SNPs. Green dots represent SNPs with a significantly different coverage between sexes based on a Wilcoxon rank sum test at the 5% significance threshold. Red dots correspond to SNPs with a significant difference in local SNPs density between sexes in a 500 SNPs region around the focal SNP based on a chi²test at a 5% significance threshold.

Finally, to focus on current intra-locus sexual conflicts rather than sex differentiation or resolved sexual conflicts, we targeted autosomal chromosomes only and implemented advanced filtering to avoid duplicated autosomal regions on sex chromosomes. Such duplications can create spurious signals of differentiation between males and females. Even when aligning the male sequences to a reference genome containing the Y chromosome, this artefact is still a potential issue. Because males and females do not have the same number of copies of each chromosome, the absence of a duplicated gene in the reference genome will result in skewed depth of sequencing between sexes for these loci (Ratio of coverage F:M of 4/3 for an X-duplicated gene and of 2/3 for a Y-duplicated gene). Sequence similarity between autosomes and sex chromosomes can also lead to reads from the sex chromosomes mapping on the autosome, generating a departure from the expected 1:1 coverage ratio between sexes. Based on these expectations, we detected 22,825 SNPs with significant coverage bias between sexes (Fig 1C). If a duplication is ancient enough, the different copies might also have diverged, which can result in a sex-specific increase in the number of SNPs. We identified and excluded 430 SNPs that showed a significant difference in the number of surrounding SNPs between sexes (in a 500 SNP window surrounding each SNP) (Fig 1C). Some of those SNPs were associated with a sex-specific increase in heterozygosity (Fig 1C), with some alleles being only present in one sex and always in individuals that are heterozygous. Altogether, after filtering, the dataset includes 1,701,083 SNPs genotyped in 49 males and 50 females.

### II Three complementary methods to identify intra-locus sexual conflicts

#### A Single-locus signal of differentiation (SNP-by-SNP)

Because of differential mortality, intra-locus sexual conflicts are expected to lead to differences in allele frequencies between males and females. We screened the three-spined stickleback genome for SNPs showing significant differentiation between sexes. For each SNP, we estimated inter-sex F_ST_ and its associated p-value and performed a Fisher’s exact test (see methods for the detail). To reduce false positive rate, a SNP was considered a candidate for intra-locus sexual conflict if it respected the three following criteria: it was in the top 1% of the Fst distribution, had a significant F_ST_ (P <= 0.001) and it was also significant with a Fisher’s exact test (P <= 0.001). This resulted in 1,478 SNPs (0.8% of total SNPs, Fig S5) that showed significant differentiation between males and females and represent putative targets of intra-locus sexual conflicts across the autosomes of the three-spined stickleback.

#### B Cumulative signals of inter-sexual differentiation (Cumulative F_ST_)

To broaden the scope of detected intra-locus sexual conflicts, we also applied a method designed to search for weak, cumulative signals of differentiation between males and females. We assessed F_ST_ significance using the null distribution for inter-sex Fst with p-values of 5%, 1% and 0.1% to call outlier, and compared the number of significant SNPs obtained to random permutation of fish from different sexes. At the genome wide scale, this method revealed no significant signals of accumulation of inter-sex differentiation at any level of F_ST_ significance (Quantile 95%: mean ratio = 0.995, 95% confidence interval = [0.991-0.998], Quantile 99% = 0.993, [0.986-0.999] and Quantile 99.9% = 0.992, [0.979-1,004], Fig S6A).At the chromosome scale, however, the pattern was clearly heterogeneous across the genome (table S2) as we detected enrichment in putative intra-locus sexual conflicts on ChrIII, ChrV, ChrVIII, ChrIX, ChrXII, ChrXIII, ChrXV, and ChrXVIII (Fig S6B).

#### C Redundancy Analyses (RDA)

We then performed a redundancy analysis (RDA) to search for multivariate associations between genotype and sex. We applied this method to identify groups of SNPs that co-vary with sex. Over the entire dataset, we found no evidence for intra-locus sexual conflicts (p-value of 0.447 and R^2^ adjusted of 1.101 ×10^−5^, table S2). Running our RDA model independently for each chromosome identified two chromosomes in which genetic variation was significantly associated with inter-sex differentiation: chrIX (p value of 0.046 and an adjusted R^2^ of 3.735 ×10^−4^) and chrXII (p value of 0.019 and an adjusted R2 of 2.271 ×10^−4^) (table S2). Both chromosomes were also significantly enriched for SNPs differentiated between males and females in the cumulative F_ST_ analysis.

### II Detecting different levels of inter-sex differentiation

The Cumulative F_ST_ and RDA approaches are strongly impacted by the scale at which they are applied, as shown by the differences between the global and chromosomal analyses. To avoid missing local effects that could be masked at the global and chromosomal scales, as well as for identifying more precisely SNPs that might be evolving under intra-locus sexual conflicts, we performed the Cumulative F_ST_ and RDA analyses in 250kb sliding windows along the genome with 50kb steps. To reduce the impact of false positives, we focused on the 1% quantile of 250kb windows with the strongest signal (highest SNP count ratio for Cumulative F_ST_, lowest p-value for RDA) for subsequent analyses. This led to 80 windows (997 SNPs) for Cumulative F_ST_, and 80 windows (174 SNPs) for RDA (Fig 2). Similarly, to reduce false positives in the SNP-by-SNP approach, we focused on the 1% 250kb windows with the highest number of significant SNPs, keeping 91 windows (192 SNPs, Fig3 A).

**Figure 2:**
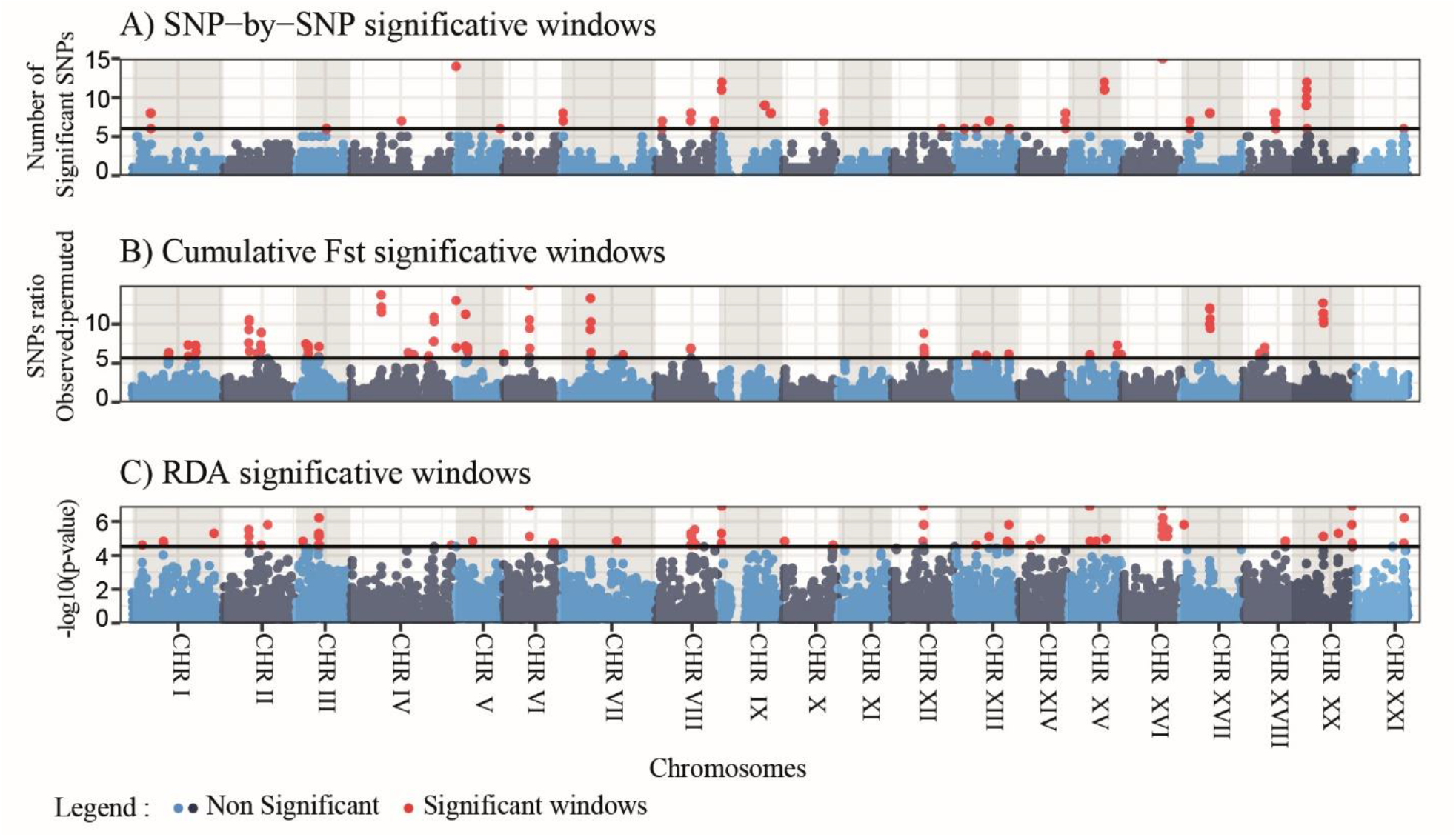
Genomic landscape of putative signatures of intra-locus sexual conflicts as detected by three approaches. Manhattan plot of significant windows using the SNP-by-SNP approach (A), the cumulative F_ST_ approach (B) and the RDA analysis (C). Red dots correspond to significant 250kb windows, and black dots represent significant windows that contain significant SNPs in linkage disequilibrium with SNPs potentially duplicated on sexual chromosomes. Other points represent non-significant windows, and the solid black line represents the significance threshold at 99%.

The three methods showed almost no overlap neither at the chromosomal scale (only chrIII, chrXII and chrXV are significant with both the Cumulative F_ST_ and RDA) nor at the window scale (0.4% and 2% of overlapping significant windows between the RDA and Cumulative F_ST_ methods and between RDA and SNP-by-SNP respectively, 3.7% between cumulative F_ST_ and SNP-by-SNP). When measuring male-female differentiation by F_ST_, we observed that Cumulative Fst and RDA detected SNPs with a significantly lower intersex F_ST_ compared to the SNP-by-SNP approach (Fig 3B), possibly providing more sensitivity to detect weak effects of sexual conflicts. Altogether, this suggests that combining these three approaches did widen the scope of sex-differentiated regions that could be detected.

**Figure 3:**
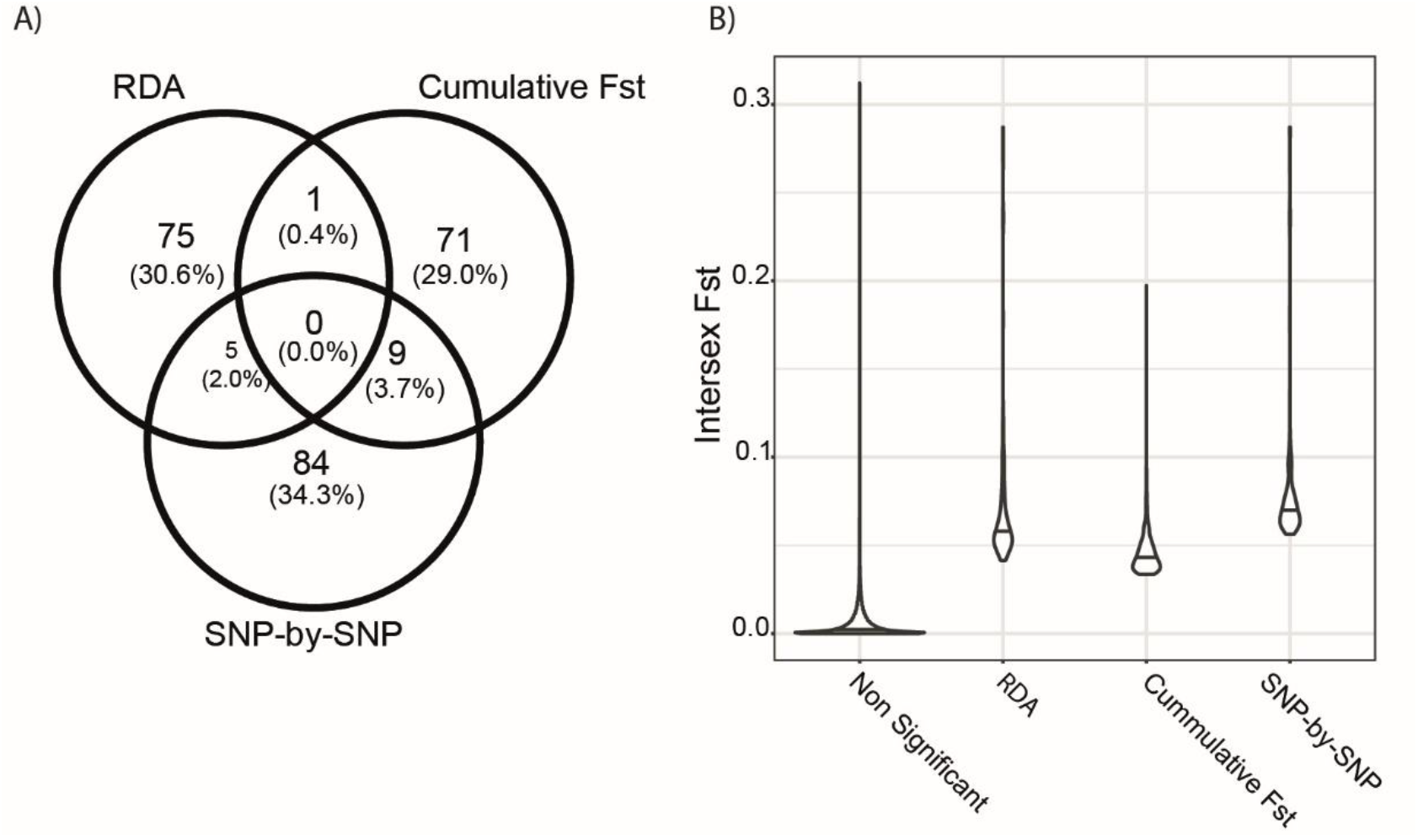
Overlap of signatures of intra-locus sexual conflicts between methods. A) Venn diagram of the overlap between significant windows for the three methods. B) Per SNP intersex Weir & Cockerham F_ST_ for significant SNPs detected by each method and nonsignificant SNPs. All pairwise comparisons are significant based on a Wilcoxon rank-sum test at a 5% significance threshold.

### IV Intra-locus sexual conflicts are enriched in functions associated with the immune system

Functional enrichment of genes within 1kb of a significant SNP showed signals for enrichment only with the Cumulative F_ST_ approach. We found significant excess of genes involved in adaptive immune response (29 out of 385 identified genes, q-value = 4.92e-09 with Benjamini-Hochberg (BH) correction (Benjamini and Hochberg 1995), as well as various processes associated with cell adhesion (*e*.*g*, cell-cell adhesion, 33 out of 385 identified genes for a q-value of 1.52e-33 after BH correction) (Table S3). We found no enrichment with the RDA and SNP-by-SNP analyses, but we also had less detection power (85 and 74 genes identified by each method, respectively).

### V The relationship between intra-locus sexual conflict and genetic diversity depends on the method of detection and is affected by recombination heterogeneity

The relationship between Tajima’s D and nucleotide diversity (π) with intra-locus sexual conflict varied depending on the detection method (Fig. 4). The SNP-by-SNP method showed an increase in π and Tajima’s D in regions associated with intra-locus sexual conflict compared to the rest of the genome (Wilcoxon rank sum test, Tajima’s D: p = 9.8×10^−3^ and π: p = 4×10^−2^). Conversely, windows identified by Cumulative F_ST_ revealed slightly lower nucleotide diversity and Tajima’s Din regions associated with intra-locus sexual conflict compared to the rest of the genome (Wilcoxon rank sum test, Tajima’s D: p = 7.54×10^−8^ and π: p = 2.6×10^−10^) while windows identified by RDA show no differences from the rest of the genome (Wilcoxon rank sum test, Tajima’s D: p = 6.3×10^−2^ and π: p = 6.8×10^−2^). However, the heterogeneity in recombination rates along the genome is also worth considering before interpretation. In fact, we noted variation in the mean recombination rates of significant regions identified with SNP-by-SNP and Cumulative Fst methods (Wilcoxon rank sum test, SNP-by-SNP: p = 2.5×10^^-2^ and Cumulative F_ST_: p = 1.3×10^^-10^ for) (Fig 4), with SNP-by-SNP being biased toward elevated recombination rates and cumulative F_ST_ toward lower recombination rates. Because recombination rate tends to be positively correlated with genetic diversity (Payseur and Nachman 2002), we performed our analyses again by comparing significant regions and other genomic regions with similar recombination rates. To do this, we divided our dataset into low (<= 1cm/Mb), medium (]1-5] cM/Mb], and high (> 5 cM/Mb) recombination rates. Doing so removed all signals of associations between genetic diversity/Tajima’s D and regions associated with intra-locus sexual conflict as detected by the SNP-by-SNP method (Fig 5). However, regions detected by RDA in medium recombination regions showed a slightly lower Tajima’s D distribution compared to the rest of the genome (p = of 2.3×10^^-3^). This was also observed for regions detected by Cumulative F_ST_ in low (p = of 4.6×10^^-2^) and medium (p-value of 4.4×10^^-2^) recombination regions. Similar observations were made for nucleotide diversity and regions detected with RDA (p = of 3.9×10^^-5^) and Cumulative F_ST_ (p = of 6.1x^10^-9^) in medium recombination regions (Fig. S7)

**Figure 4.**
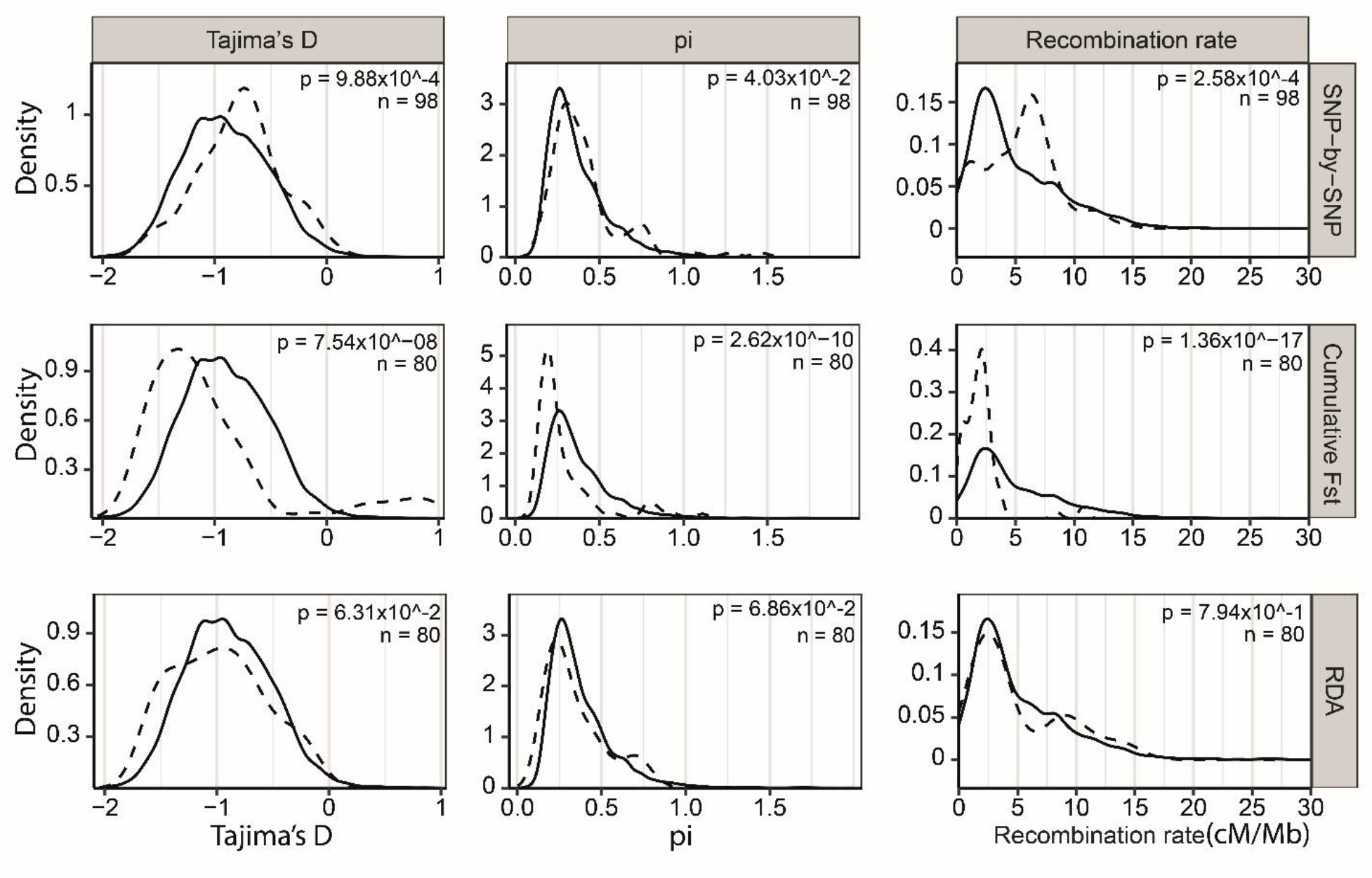
Association between signatures of putative intra-locus sexual conflicts and genetic metrics. Distribution of Tajima’s D, π, and recombination rates (from left to right) for windows detected by each method (dashed line) as significantly associated with sexual conflict compared to the bulk of the genome (solid line); From top to bottom, SNP-by-SNP, RDA and: Cumulative F_ST_. p represents the P-value of a Wilcoxon rank-sum test comparing the distribution of significant and non-significant windows; n represents the number of significant windows.

**Figure 5:**
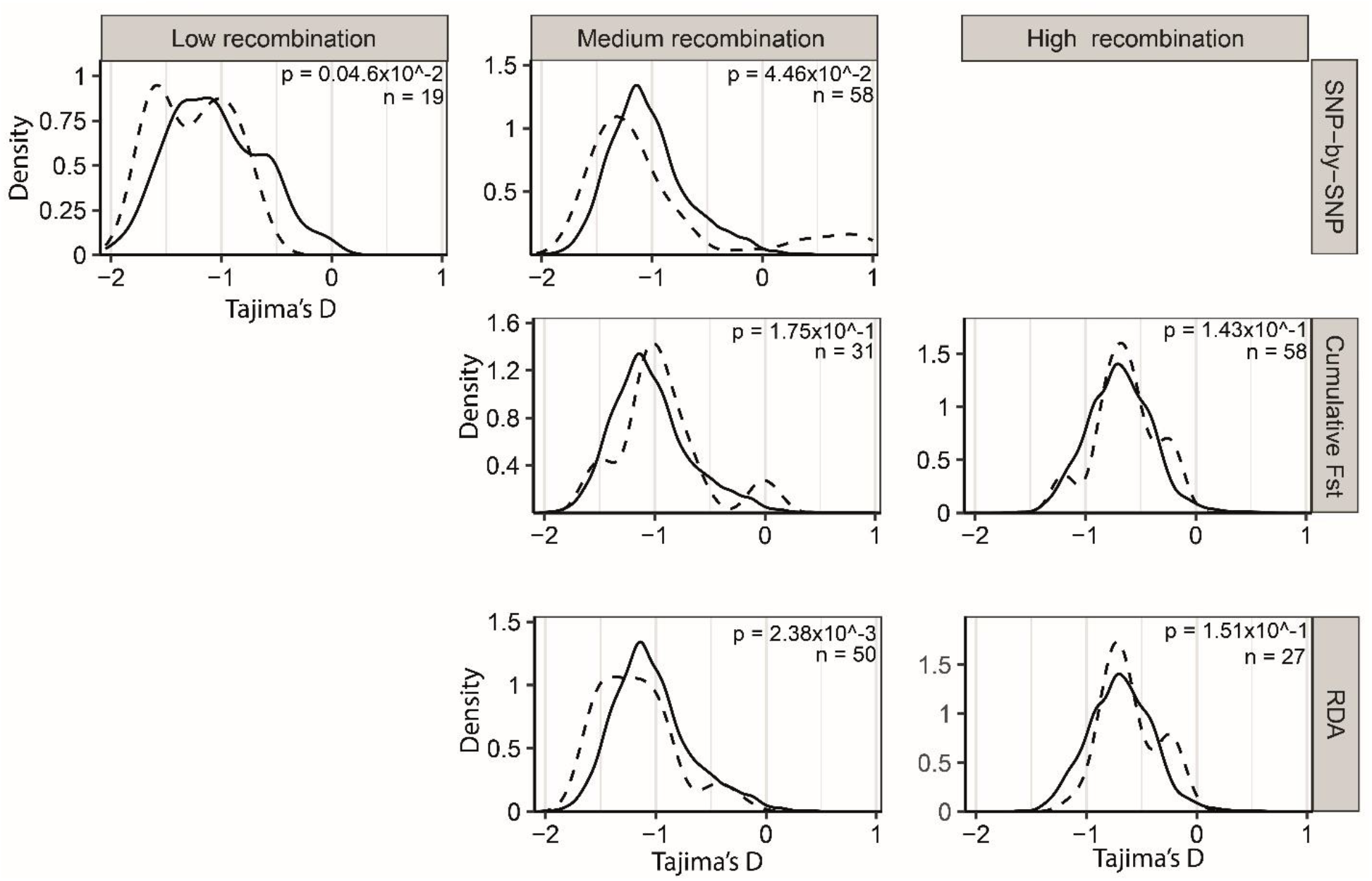
Association between putative intra-locus sexual conflict and Tajima’s D when controlling for recombination rate. Distribution of Tajima’s D for each detection method (dashed line) compared to the bulk of the genome (solid line) for each category of recombination rate (low, medium, high, from left to right). From top to bottom: Cumulative F_STt_, RDA and SNP-by-SNP. For each method, only category of recombination rate represented by at least 10 windows associated with putative intralocus sexual conflict are represented; p represents the p-value of a Wilcoxon rank-sum test comparing the distribution of significant and non-significant windows; n represents the number of significant windows.

### V1 Rare intra-locus sexual conflicts are associated with increased genetic diversity

We found four genomic regions associated with intra-locus sexual conflict that showed potential signals of long-term balancing selection (Tajima’s D > 0), Fig. 6). Two are located on chrIV ([9.6Mb-10.45Mb] and [20.15Mb – 20.9 Mb]) and two on chrVII ([0.2Mb - 1.05Mb], [9.15Mb – 10.15Mb]). Significant SNPs in those regions were still associated with elevated Tajima’s D values at a finer scale (25kb sliding windows). Pairwise estimation of r²between all pairs of SNPs in those regions showed that significant SNPs were associated with elevated linkage disequilibrium in two regions on chrIV and one region on chrVII ([9.15Mb – 10.15Mb]).

**Figure 6:**
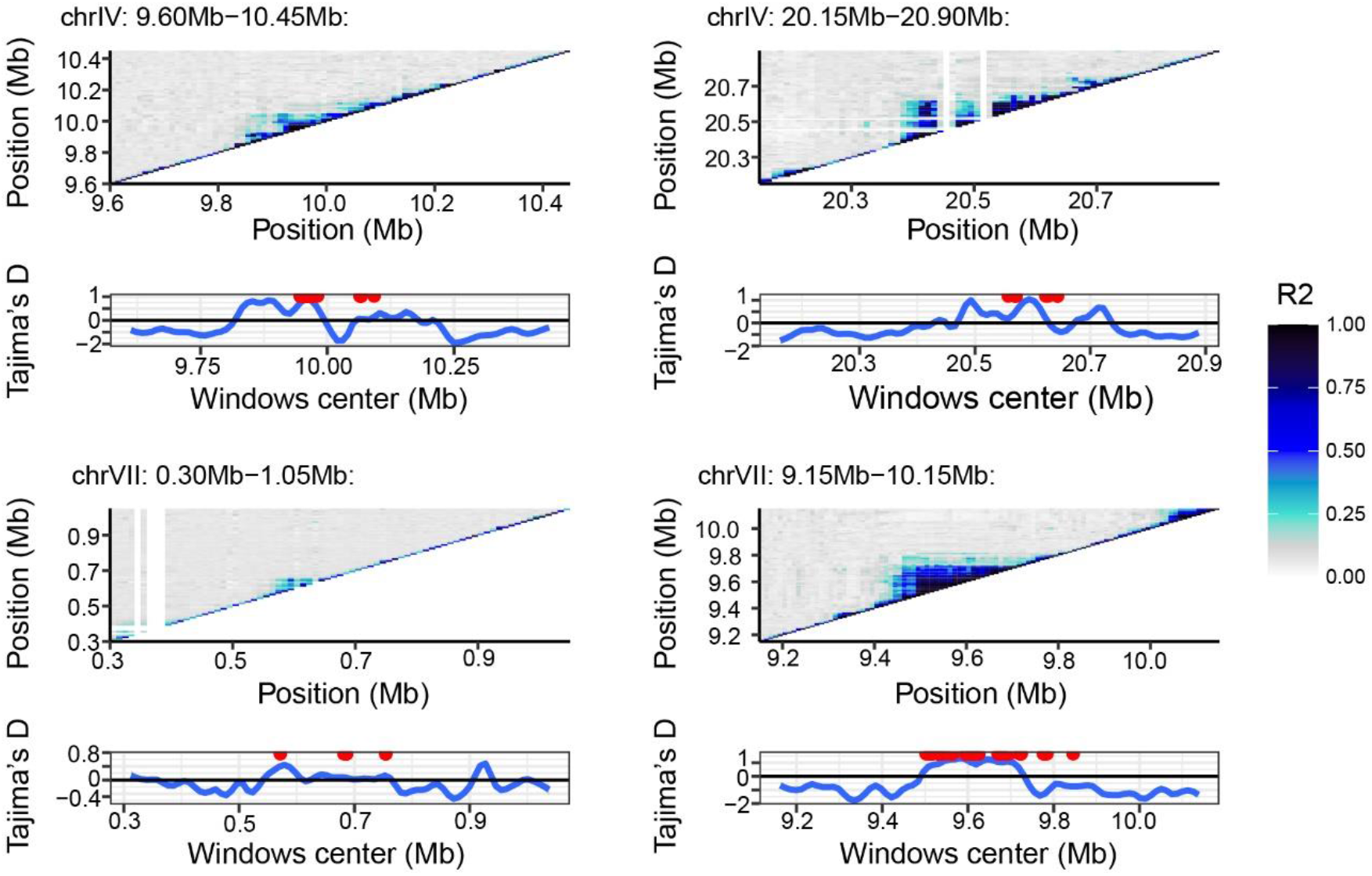
Zoom in on regions associated with elevated Tajima’s D. A) Pairwise linkage disequilibrium in regions significantly associated with putative intra-locus sexual conflict (extended by 250kb in each direction for visibility) and with a positive Tajima’s D, averaged over 10kb windows. B) Tajima’s D estimated in 25kb windows. Red dots indicate the location of SNPs associated with the signal of intra-locus sexual conflict.

We found 21 significant SNPs in regions that are within 1kb or less of an annotated gene, resulting in seven genes associated with intra-locus sexual conflicts and balancing selection (table 1). On these seven genes, three are associated with immune functions or show partial homology to immunoglobulin (*dock11, ENSGACG00000019964, ENSGACG00000019965*). One gene shows similarity to genes involved in cellular processes (*ENSGACG00000018638*). *KIRREL1a* is associated with cell-cell junction. *ANO4* is associated with a transport function and has been found to be potentially associated with adaptation to altitude in the Tibetan chicken (Zhang et al. 2016). We were unable to associate the gene *maker-chrVII-exonerate_protein2genome-gene-7*.*17-mRNA-1* to any function, GO or clear blastx results. When checking for coverage bias between sexes that could show partial duplication or similarity with sex chromosomes, only *ENSGACG00000019965* showed a departure from the 1:1 expected ratio compared to surrounding noise, with an increased coverage in males compared to females (Fig. S8)

**Table 1:**
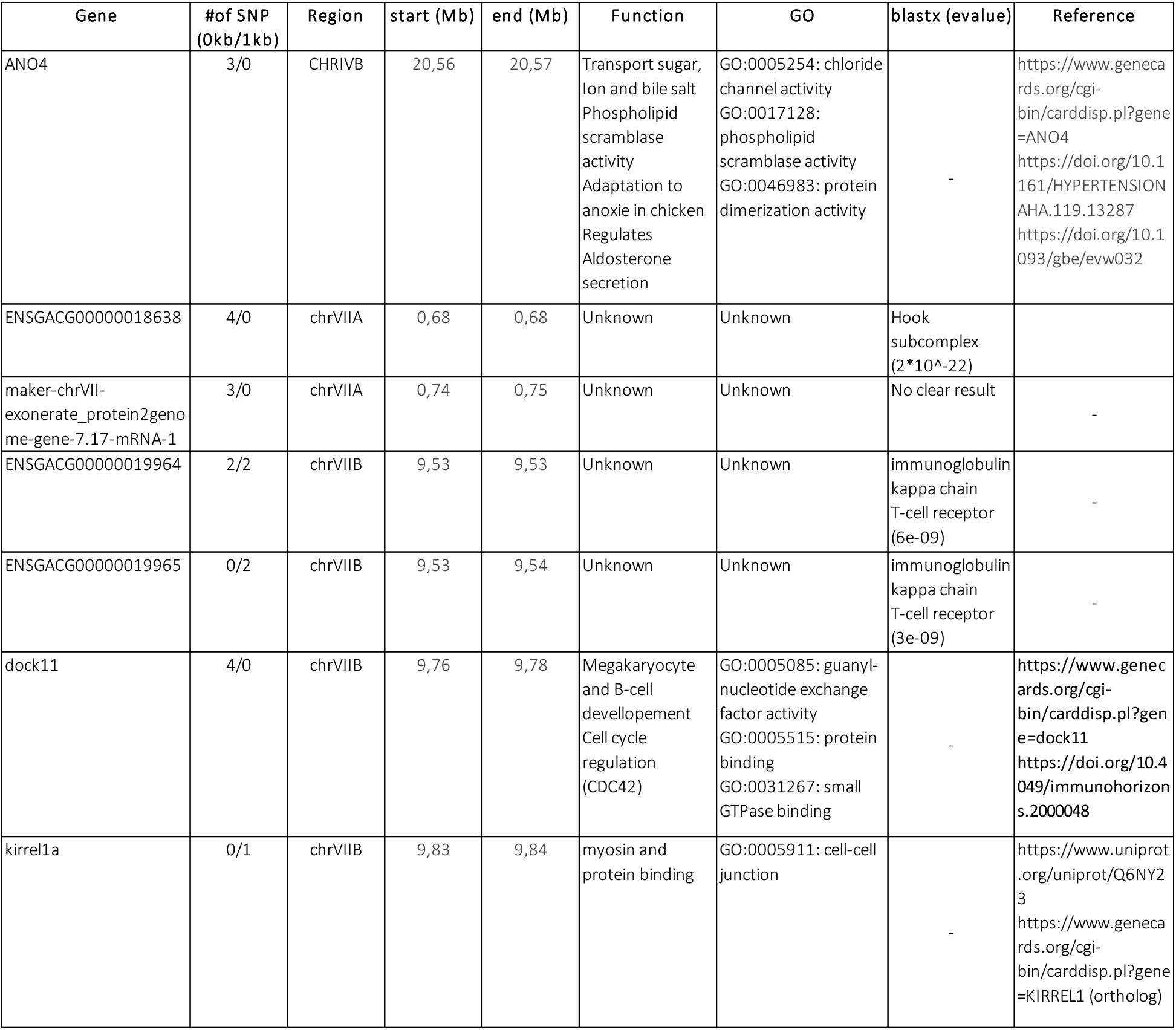
Summary of genes associated with intra-locus sexual conflict in regions under potential balancing selection. A and B letters are used to differentiate different regions of potential balancing selection on the same chromosome

## Discussion

Using whole genome resequencing of 49 male and 50 female three-spined stickleback, we detected SNPs and regions with signature of putative intra-locus sexual conflicts. Following stringent filtering of genomic regions duplicated on sex chromosomes, we combined three methods (SNP-by-SNP, Cumulative F_ST_ and RDA) to detect signals of inter-sex differentiation. Focusing on the 240 windows with the strongest signals, we showed that most intra-locus sexual conflicts were not associated with increased genetic diversity. However, we still identified four genomic regions that were associated with signals of balancing selection.

### I–Sexual chromosomes as hotspot of sexual conflict resolution

Our data revealed putative duplications between sex chromosomes and autosomes, which affected 1.35 % of SNPs (n=23,282). Such a signal has often been attributed to the lack of Y reference, which leads to reads belonging to two different copies of a duplicated sequence mapping at a single position on an autosome. For the three-spined stickleback, we relied on a high-quality reference of the Y chromosome but still found numerous duplicated regions (56.4% of all potentially duplicated SNPs are associated with the Y chromosome, Fig. 1B). These duplicated SNPs are likely to be caused either by high similarity between autosomal and sex-linked copies or by duplications that are not represented in the Y sequence, such as duplications that are specific to our population or that are difficult to assemble. This pattern is consistent with previous studies in the trinidadian guppy (*Poecilia reticulata*) and three-spined stickleback (Bissegger et al. 2020; Lin et al. 2022). We also identified duplications between autosomes and the X chromosome (43.3 % of all potentially duplicated SNPs) that were caused by the same artefact. Moreover, we found that potentially duplicated X-chromosome SNPs (i.e., those showing excess of coverage or polymorphism in females) often exhibit increased heterozygosity in females only. However, duplication on the X chromosome would also lead to an increase in male heterozygosity, even if at a lower level, and therefore probably does not explain the observed pattern. Those regions could be duplicated on both the X and the Y chromosomes but correctly assembled on the Y chromosome only, resulting in reads from the X chromosomes mapping on their autosomal copy in females but in their Y copy in males. This would create a female-specific increase in coverage and heterozygosity and would not happen in genomes missing a reference for the Y chromosomes. We believe this is a more likely explanation of the observed heterozygosity bias.

Artefactual polymorphism generated by duplicated genomic regions that are collapsed in the assembly represents an analytical challenge for population genomics and this matter is increasingly recognized in different biological systems. For instance, in the American lobster (*Homarus americanus*), SNPs associated with geographical variation also show variation in heterozygosity and relative coverage of the reference and alternative alleles (Dorant et al. 2020). In selfing *Arabidopsis thaliana*, numerous SNPs with unexpected heterozygosity levels were in fact caused by duplicated regions of the genome that were missing from the reference (Jaegle et al. 2022). Observed excess of heterozygosity then occurs when reads originating from two diverged copies align at a single locus. This is an issue with any genome alignment-based method and must be addressed through proper filtration. However, it can lead to more complex biases that are difficult to correct for when duplication patterns are sex-linked. This observation reinforces the idea that caution is needed when analyzing and interpreting large genomic datasets containing both males and females.

Duplications between autosomes and sex chromosomes can also contribute to resolve sexual conflicts (Mank et al. 2020; Lin et al. 2022) and as such, are biologically meaningful. Similar to the recent study of Bissegger et al. (2020b) on three-spined stickleback, we identified substantial duplications on the Y chromosome, which is expected to retain gene copies and alleles that are advantageous to males (Bachtrog 2013; Carvalho et al. 2015) and be a hotspot for conflict resolution. While we did not directly test this hypothesis, these duplications might be remnants of resolved intra-locus sexual conflicts. We did, however, identify more complex patterns of X and X-Y potential duplications, which, interpreted in the same context, suggests that the X chromosome might also play a role in the resolution of intra-locus sexual conflicts. Altogether, these results illustrate a dynamic process of duplicating parts of the autosomes into the sex chromosomes. Further research focussing on the timing of gene duplication from autosomes to sex chromosomes and comparing these dynamics in populations with a gradient of differentiation would help clarifying the role of intra-locus sexual conflict in the divergence of X and Y chromosomes.

### II Signature of intra-locus sexual conflict as detected by a complementary set of methods

Through differential mortality, intra-locus sexual conflicts are predicted to translate into differential allelic frequency between sexes (Cheng and Kirkpatrick 2016; Lucotte et al. 2016; Lin et al. 2022), which can be detected using various approaches such as inter-sex F_ST_ (Wright et al. 2018; Lin et al. 2022) or genome-wide association studies (GWAS, (Ruzicka et al. 2020; Kasimatis et al. 2021). However, substantial mortality is required to generate significant inter-sex differentiation and statistical power to detect this signal is low, especially after accounting for multiple testing in genome-wide datasets. Since many loci under strong intra-locus sexual conflict over mortality would likely represent an excessive mortality cost for natural populations, the search for intra-locus sexual conflicts in genome-wide datasets mainly focus on finding small-effect variants. This may be done by reducing the false discovery rate corrections at the gene level (Lucotte et al. 2016) or by focusing on genes showing cumulative signals of inter-sex differentiation (Ruzicka et al. 2020; Lin et al. 2022). Adapting these approaches to non-coding regions (SNP-by-SNP and Cumulative F_ST,_ Fig. S5, S6) and using multivariate approach (RDA) to detect potential patterns of polygenic selection (table S2) in the three-spined stickleback, a species with large juvenile cohorts (Whoriskey et al. 1986) and with documented differences in predation level between sexes at the adult stage (Whoriskey and Fitzgerald 1985), revealed very limited signals of intra-locus sexual conflict. These results are in line with those of Lin et al. (2022) who found limited evidence for intra-locus sexual conflict in the Trinidadian guppy, which display sex-specific susceptibility to mortality.

When looking at more specific targets of intra-locus sexual conflict in the genome by applying our methods to 250kb sliding windows and focusing on the strongest signals, we found that our three methods detected different degrees of per-SNP male-female differentiation (Fig 3B) and that there is very little overlap between them (Fig. 4A). In particular, the RDA and Cumulative F_ST_ methods were able to detect weaker signals of inter-sex differentiation than the SNP-by-SNP approach, allowing the detection of smaller effect variants than the SNP-by-SNP approach (Fig. 4 B), although they were sensitive to the scale at which they were applied. Importantly, the Cumulative Fst method represents a measure of the proportion of true positives among the significant SNPs in a given genomic region, but cannot distinguish between true and false positives (Ruzicka et al. 2020). this method thus complicates the identification of precise targets of intra-locus sexual conflict. Using complementary methods effectively allowed us to widen the range of detected intra-locus sexual conflicts and can improve our ability to document such conflicts.

### III Intra-locus Sexual Conflicts are not Associated with Increased Genetic Diversity

Theory predicts that intra-locus sexual conflicts could locally increase genetic diversity. This prediction is sometimes taken for granted and some studies have used signature of increased genetic diversity, such as elevated Tajima’s D, to confirm the identification of intra-locus sexual conflict (Wright et al. 2018; Wright et al. 2019). However, the extent to which this relationship is valid across the genome is still lacking. Studies focusing on specific traits or genes have shown that intra-locus sexual conflict can maintain genetic diversity in a variety of context (Foerster et al. 2007; Barson et al. 2015; Hawkes et al. 2016; Lonn et al. 2017), but only three studies have tested this relationship directly in large genomic datasets (Dutoit et al. 2018; Sayadi et al. 2019; Lin et al. 2022). These studies led to variable conclusions. Dutoit et al. (2018) found a positive association between signatures of intra-locus sexual conflict and genetic diversity, while Lin et al. (2022) did not and results from Sayadu et al. (2019) suggested that intra-locus sexual conflict might maintain genetic diversity through relaxed purifying selection. One explanation for this discrepancy may be that Dutoit et al. (2018) did not account for the effect of sex-chromosome duplications, which can create regions of artificially elevated genetic diversity mistaken for signal of balancing selection. Our results are more in line with those of Lin et al. (Lin et al. 2022) as we did not find global associations between signatures of intra-locus sexual conflict and increased genetic diversity measured by Tajima`s D and nucleotidic diversity (Fig. 5).

Tajima’s D and nucleotide diversity are metrics that usually detect medium-to-long term excess in polymorphism (Fijarczyk and Babik 2015). Hence, we may be missing signals of recent balancing selection or locally relaxed purifying selection instead of long-term balancing selection similar to those identified by Sayadi et al. (2019). These results support the hypothesis that most intra-locus sexual conflicts could be either resolved over short evolutionary time scales or that they are not stable over time. Resolution of intra-locus sexual conflict implies decoupling the genetic bases of the concerned traits between males and females. This can be achieved through gene duplications on sex chromosomes, as discussed before, but also through other processes that can modulate the effect of a genotype in a sex specific way (van der Bijl and Mank 2021). In sticklebacks, for instance, sex-biased patterns of gene expression or alternative splicing have been documented (Naftaly et al. 2021; Kaitetzidou et al. 2022).

However, the conditions for intra-locus sexual conflict stability depend on the strength of selection, with conditions for stability narrowing when selective coefficients are lower (Zajitschek and Connallon 2018). As we mainly detected variants with small inter-sex differentiation, it is very likely that a combination of dynamic resolution of intra-locus sexual conflicts and instability is one reason why we observed only four regions under potential conflicts associated with balancing selection.

Because our identification of intra-locus sexual conflict was based on significant differences in allelic frequency between sexes, directional selection in one sex could also be detected by our approach. In that case, we would not expect an increase in genetic diversity around the selected locus, and such loci would dilute observed genetic diversity maintained at the identified loci. This is supported by our finding that genetic diversity was slightly lower around putative loci under sexual conflict in regions of medium recombination rate for RDA and Cumulative F_ST_ windows (Fig 5, S7), which is concordant with local directional selection in one sex leading to a soft selective sweep. These results confirm that methods classically used to detect sex differences may likely detect a combination of sex-specific selection and intra-locus sexual conflict. Monitoring variation in allelic frequency in each sex across age classes or direct measurement of survival variation according to sex and genotype are needed to segregate the effects of intra-locus sexual conflict and sex-specific selection over survival.

The SNP-by-SNP method revealed that regions putatively involved in intra-locus sexual conflict were located in genomic regions of higher recombination rates relative to the rest of the genome. This might be explained by method-specific sensitivity. Indeed, because the SNP-by-SNP approach is based on a fixed number of significant SNPs in a given genomic region, regions with higher recombination rates leading to an increase in genetic diversity are more likely to be significant than regions with lower recombination rates (Payseur and Nachman 2002). Similarly, lower recombination regions lead to higher linkage disequilibrium, meaning that an intra-locus sexual conflict could generates a signal of intersex differentiation to a larger neutral region around the causal SNP(s). At the opposite, when permuting our dataset with the Cumulative F_ST_ method, permutations would result in losing signal at more SNPs for the same reason, leading to the Cumulative F_ST_ method being more powerful in lower recombination regions. This emphasizes the need for comparing, when possible, regions of similar recombination rates when studying genetic diversity. However, by splitting our dataset by the level of recombination, we might also lack the power to detect subtle increases in genetic diversity.

### IV Rare intra-locus sexual conflicts are associated with increased genetic diversity

Four genomic regions with sexual conflict (two on chrIV, two on chr VII and one on chrXXI) showed elevated genetic diversity (Tajima’s D >=0), three of which also exhibited increased linkage disequilibrium (Fig. 6). Both of these statistics can be indicative of balancing selection (Fijarczyk and Babik 2015). In three-spined stickleback, balancing selection has previously been reported to be occurring mainly on chrIV, chrVII and chrXXI (Thorburn et al. 2021). These chromosomes have also been shown to be involved in controlling the expression of many phenotypic traits (Peichel and Marques 2017; Rennison et al. 2019; Thorburn et al. 2021) that exhibit differences between sexes, *e*.*g*, pigmentation, body size, reproduction (for chrIV and chrXXI) and defence (for chrVII, measured as dorsal spine and lateral plate morphology) (Craig and FitzGerald 1982; Poulin and Fitzgerald 1989; Kitano et al. 2007, Reimchen and Nosil 2001). Those results are concordant with our interpretation that these regions of increased genetic diversity could be evolving under balancing selection driven by intra-locus sexual conflicts.

*ANO4* is associated with various biological processes, including aldosterone secretion, which appears to be associated with osmoregulation in three-spined stickleback (Li and Kültz 2020). *ANO4* has also been found in association with adaptation to altitude in Tibetan chicken (Zhang et al. 2016). In our population, three-spined sticklebacks reproduce in tide pools with very low dissolved O2 and high salinity (Whoriskey et al. 1986; Poulin and Fitzgerald 1989, personal observations). Given the higher energetic cost for males during the reproduction period (Chellappa and Huntingford 1989; Dufresne et al. 1990), and decreased tolerance to salinity during reproductive period (Wootton 1984), the occurrence of intra-locus sexual conflicts over potential adaptation to low O2 concentration or elevated salinity represents a plausible hypothesis. *Dock11* is associated with functions of the immune system (Sugiyama et al. 2022, and see its GeneCards entry, Stelzer et al. 2016), and we found that *ENSGACG00000019964* and *ENSGACG00000019965* show similarity with immunoglobulin (Table 1). *ENSGACG00000019965* shows patterns of male-bias coverage on its larger exon, suggesting potential duplication on the Y chromosome (Fig. S8). Also, different parasite species and abundance has been documented between sexes in three-spined stickleback (Reimchen and Nosil 2001), even though they were not necessarily associated with conflict over immunity (De Lisle and Bolnick 2021). Moreover, the immune system can play a role in sexual selection (Folstad et al. 1994) and affect variation in morphology (De Lisle and Bolnick 2021), suggesting that mechanisms other than intra-locus sexual conflict could explain the differences observed between sexes. However, a study in *Drosophila melanogaster* suggests that intra-locus sexual conflict might also be a driver of immune system evolution (Morrow and Innocenti 2012) and genes of the immune system are often associated with balancing selection (Andres et al. 2009). Coupled with the GO enrichment in immune system function observed with the Cumulative F_ST_ method, it is likely that the interplay between the intra-locus sexual conflicts dynamics and the context of the immune system favour the maintenance of genetic diversity in those regions.

## Conclusion

The combination of three complementary detection methods (SNP-by-SNP, Cumulative F_ST_ and RDA) revealed modest yet significant signatures of intra-locus sexual conflict of various strengths throughout the three-spined stickleback genome. These were mainly associated with immunological and neuronal development processes. We found that most intra-locus sexual conflicts do not drive long-term balancing selection or increased genetic diversity, suggesting that they are a dynamic process and are either quickly resolved or that selective pressures are varying through time. Those conflicts may however still play a role in maintaining genetic diversity over shorter timescales by locally reducing the effect of background selection. We also showed that false positives can arise from the duplication of genomic regions into the sex chromosomes, highlighting the importance of accounting for group-specific rearrangements in population genomic studies.

## Methods

We collected 100 adult anadromous three-spined sticklebacks between May and July 2018, at Baie de l’Isle verte, Québec (48.009961, -69.407070), which belong to a single panmictic population (McCairns and Bernatchez 2009). To limit the potential impact of environmental heterogeneity and fine scale genetic structure, we sampled all individuals in the same tide pool. Fins were preserved in 95-98% ethanol and we extracted DNA following a salt extraction protocol modified from Aljanabi and Martinez **(1997**). We checked DNA quality on a 1% agarose gel, and measured DNA concentration and contamination using a Thermo Scientific™NanoDrop™2000 spectrophotometer and the Biotium AccuClear Ultra High Sensitivity dsDNA Quantitation Kit. Samples were sent the Centre d’Expertise et de Services Genome Québec (Montréal, QC Canada). for NEB Ultra II Shotgun DNA library preparation and quality checking, followed by whole genome sequencing on nine illumina HiseqX lanes with a targeted coverage of 15X.

### Genome Alignment and Cleaning

We first ran fastsp version 0.15.0 (Chen et al. 2018) with default settings on raw reads to trim Illumina adapters and filter for quality. We used bwa-mem version 0.7.17-r1188 (Li 2013) to map the sequencing reads to the 5^th^ version of the stickleback reference genome (Nath et al. 2021) using a sorted alignment with samtools version 1.8 (Li et al. 2009). Males and females were respectively mapped on a reference including or not the Y chromosome reference (Peichel et al. 2020), available on https://stickleback.genetics.ugq.edu). The Y chromosome was trimmed of its pseudo-autosomal region (first 2.5Mb) as it was already present on the X reference (chrXIX). We then used picardstools MarkDuplicates (1.119, http://broadinstitute.github.io/picard/) to identify and trim PCR duplicates, and GATK RealignerTargetCreator and IndelRealiner (DePristo et al. 2011) to perform local realignment around putative indels and to improve the read mapping in those areas. Finally, we clipped overlapping read pairs using bamUtils clipOverlap version 1.0.14 (Jun et al. 2015) and removed reads that became unmapped in the process using samtools view (-F 4 option). Beforehand, we used bwa index and picardstools CreatesSequenceDictionary to build requirements for reference genome handling by the various tools. We used samtools index to index bam files between each step after alignment, except before indel realignment, which required using picardstools BuildBamIndex.

### Variant Calling

We used bcftools v1.12 to call autosomal SNPs using mpileup (-a AD,DP,SP,ADF,ADR, -q 5, -d 20000) coupled with bcftools calls (Li 2011) (-a GP, GQ, -G -). Resulting biallelic SNPs were filtered using bcftools based on several metrics: 1) Mapping quality (>= 30), Mapping quality bias (p > 10^-3), Read position bias (p > 10^-3), and QUAL >= 25. We also used custom python3 scripts to calculate Strand Odds Ratio as suggested by GATK (https://gatk.broadinstitute.org/hc/en-us/articles/360036361772-StrandOddsRatio) and removed all SNPs with a value above 3.5. We changed all genotypes with coverage <4 (to avoid genotype mis-calling) or > 35 (to eliminate SNPs in duplicated or repetitive regions) to missing data and removed SNPs with more than 80% missing data. We used vcftools v1.1.16 (Danecek et al. 2011) to exclude individuals with more than 10% missing data, and ran the process again on the remaining individuals. Because rare variants are unlikely to be significant (because of power limitation) in our dataset, we removed SNPs with minor allele frequency less than 10%. Unless mentioned otherwise, the following analyses were performed using R 4.0.3 (R Core Team 2014) and python version 3.7.4 (Van Rossum and Drake 2009).

### Detection of large structural variants

Because large inversions are known to play a role in three-spined sticklebacks’ evolution (e.g., adaptation to marine / freshwater (Jones et al. 2012; Liu et al. 2018) and can be under strong selection as they form long haplotypes involving several genes, we scanned the genome to identify putative polymorphic inversions in our population. To detect putative polymorphic chromosomal rearrangements, we used the lostruct R package (Li and Ralph 2019, available at https://github.com/petrelharp/local_pca) to perform a local Principal Component Analyses (PCA) in 500 SNP sliding windows across the genome. We then performed a Non-Metric Multidimensional scaling (NMDS) analysis on these PCAs to delimit genomic regions in which PCAs are highly correlated, which is a typical signature of large non-recombining rearrangements. Those regions were excluded from the main dataset and analysed separately.

To further characterize these regions, we performed a PCA using the prcomp function from the stats R package on each of the identified regions and used a k-means analysis to assign individuals to putative clusters. For all PCAs and for the redundancy analysis (RDA, see below), missing data were imputed with the most frequent genotype and data were centered but not scaled. For putative inversion, we typically expect to identify three groups corresponding two the two homozygous haplogroups and heterozygotes. We performed a chi²test to test for Hardy Weinberg equilibrium of the putative variants. We then estimated variation in heterozygosity and mean depth of coverage along the putative structural variants for each genotypic group present on the PCA. We also tested for potential ongoing intra-locus sexual conflicts using a Fisher’s exact test of the sex ratio among the different groups identified by PCA, as well as inter-sex F_ST_ within each group on the PCA and all groups confounded.

### Population structure

After removing large genomic rearrangements, we searched for potential sex-specific structure that might lead to significant differences between males and females. We used two approaches to search for atypical groupings of individuals. First, we imputed our dataset, replacing missing data by the most frequent genotype, and then ran a PCA to search for axes segregating males and females. Second, we compared the relatedness among males and females using the implementation of the KING equation to infer pairwise relationship coefficients among individuals (Manichaikul et al. 2010).

### Duplications on sex chromosomes

Autosomal regions duplicated on sex chromosomes can create spurious signals of differentiation between males and females. Recent work revealed that aligning sequencing data to a reference genome missing the hemizygous chromosome will result in reads from gene copies on the hemizygous chromosome incorrectly mapping on the autosomal copy of the gene (Kasimatis et al. 2019; Bissegger et al. 2020; Lin et al. 2022). This can create sex specific polymorphism and heterozygosity that can be mistaken for intralocus sexual conflict. Even if we did align male sequencing product to a reference genome containing the Y chromosome, this artefact is still a potential issue, as some duplicated genes can be mis-assembled in the reference genome. Because the phylogeographic origin of our northwest Atlantic population differs from that from which the reference genome was generated (a freshwater population from British Columbia, (Peichel et al. 2020; Nath et al. 2021), such duplicated regions could also be specific to our population and thus missing from the reference, leading to the same issue. Moreover, the same logic applies to the X chromosome and to more complex scenarios of gene duplications.

Here we used two methods adapted from Lin et al. (2022) to detect and filter SNPs potentially duplicated on sex chromosomes. Because males and females do not have the same number of copies of the sex chromosomes (males are heterogametic), the absence of a duplicated gene in the reference genome will result in skewed depth of sequencing between sexes (the expected ratio of coverage Females:Males is 4:3 for an X-duplicated gene and 2:3 for a Y-duplicated gene). We used a Wilcoxon rank-sum test to compare median coverage in males and females to detect SNPs with significant differences in coverage between sexes at a 5% significance threshold. If the duplication is ancient enough, the different copies might have accumulated divergence, which can result in a sex-specific difference in the number of SNPs. To detect those, we estimated the number of polymorphic SNPs in males and females in a 500-SNP window around each SNP and used a Fisher exact test to remove any SNPs with a significant difference in the number of surrounding polymorphic SNPs between males and females at a 5% significance threshold. Because we wanted to remove as many suspicious SNPs as possible rather than identifying SNPs duplicated with high confidence, we did not apply a multiple test correction.

### Detecting “true” signals of intra-locus sexual conflict

Intra-locus sexual conflicts over survival are expected to lead to differences in allele frequencies between males and females. Because we expect weak levels of inter-sex differentiation, finding the right balance between discarding false positives and maintaining stringency is a complex task. Method of false discovery rate correction on genomic datasets would require intense sampling to reach sufficient power to detect significant signals of inter-sex differentiation even on medium effect SNPs (Lucotte et al. 2016; Kasimatis et al. 2019). To identify a wide range of potential intra-locus sexual conflict across the three-spined stickleback genome, we combined three approaches with a priori different sensitivities: SNP-by-SNP, Cumulative F_ST_ and multivariate Redundancy analysis (RDA). For the first method (SNP-by-SNP), we adapted an approach developed in Lin et al. (2022). For each SNP, we estimated intersex F_ST_, as defined in Cheng and Kirkpatrick (2016), where 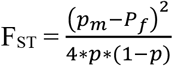where p_m_and p_f_ represent the estimated allelic frequency at a biallelic locus in males, females, respectively and p is the average of p_m_ and p_f._ We assessed its significance based on a null distribution of inter-sex F_ST_ developed in Ruzicka et al. (2020). A SNP was considered as a candidate for intra-locus sexual conflict if it was in the top 1% of the F_ST_ distribution, had a significant inter-sex F_ST_ and was significant in a Fisher’s exact test of allelic distribution between males and females, using a significance threshold of 0.001 for both tests. To further reduce false positives, we calculated the number of significant SNPs within 250kb sliding windows (50 kb steps) across the genome and focused on the 1% of windows with the most differentiated SNPs for gene enrichment analysis and to investigate the relationship between intra-locus sexual conflict and genetic diversity.

To broaden the scope of intra-locus sexual conflict detection, we also applied a method designed to search for weak, cumulative signal of differentiation between males and females (Cumulative F_ST_). We adapted a method recently proposed by Ruzicka et al. (2020), who developed a null model for inter-sex F_ST_ in the absence of natural selection and suggested looking for an excess of SNPs with significant F_ST_ between the true groups compared to the random expectation between permuted groups. We generated 100 datasets in which we permuted male and female labels and for each permutation, we estimated the ratio of the number of significant SNPs in our real dataset to the permuted datasets, considering three levels of significance: p= 0.05; 0.01; 0.001. We applied this approach at three different levels in our dataset (whole genome, by chromosome, and in local 250kb windows) to limit the effect of regions with lower-than-expected inter-sex divergence, which could reduce and cancel the signal of accumulation of significant SNPs in other regions. For the window approach, we only considered the 1% significance threshold to identify significant SNPs in each window.

Finally, Redundancy analyses (RDA) are multivariate ordination methods that have been successfully employed in genotype-environment association studies (e.g. Laporte et al. 2016; Capblancq and Forester 2021). This method works by analyzing the covariance of loci (response variable matrix) in response to environmental variation (environmental matrix) and allows identifying weak signatures of polygenic selection. In the case of intra-locus sexual conflict, it can be applied using genotypes as the response variable matrix and the sex of samples as the “environmental matrix”. Again, we considered three levels of significance (p = 0.05, 0.01, 0.001) and considered SNPs with a contribution to the constrained axis that is 2, 2.5 or 3 standard deviations (sd) from the mean as outliers. We used a similar approach as with the Cumulative F_ST_ method by running the RDA at three genomic scales: whole genome, per chromosome and in 250kb overlapping sliding windows (50kb overlap). Only the 2.5 sd significance threshold was considered for the window approach. For the Cumulative F_ST_ and RDA methods, we considered their outcomes to reflect an enrichment in intra-locus sexual conflicts if the 95% confidence interval of the mean ratio of the number of significant SNPs in the real dataset compared to random permutations did not include zero. We considered the 1% of sliding windows with the highest ratios as candidate regions for intra-locus sexual conflicts.

### Comparison of sensitivity of the methods

To determine whether the three methods we used detected regions with different levels of inter-sex differentiation, we used a Wilcoxon rank-sum test to compare the distribution of Weir & Cockerham’s q F_ST_ estimator (Weir and Cockerham 1984) as implemented in vcftools v1.1.16 for SNPs that were significant in windows identified by each of the three methods SNP-by-SNP, cumulative F_ST_ and RDA) as well as nonsignificant windows.

### GO enrichment

To test if certain biological functions were associated with intra-locus sexual conflict, we performed a Gene Ontology (GO) analysis on genes located within in 1kb of significant SNPs from significant 250kb windows in our dataset. The 1kb distance was defined according to Kratochwil and Meyer (2015) as being in regions likely to contain cis-regulatory elements. Because duplication on sex chromosomes is a way of resolving intra-locus sexual conflict, we also performed GO enrichment analysis on SNPs identified as potentially duplicated on sex chromosomes to see if we could identify similar biological processes between our putative unresolved and resolved conflicts. To do so, we annotated a three-spined stickleback transcriptome from Jones et al. (2012) using the swissprot database and downloaded gene information from the uniprot database. We then used goatools (Klopfenstein et al 2018) to perform Fisher’s exact tests and assess enrichment for biological processes using a 5% q-value threshold after a Benjamini-Hotchberg correction for false discovery rate (Benjamini and Hochberg 1995). All scripts are available at https://github.com/enormandeau/go_enrichment.

### Relationship Between Intra-Locus Sexual Conflicts and Genetic Diversity

To test if intra-locus sexual conflicts were associated with increased genetic diversity across the genome, we estimated two classically used metrics; the Tajima’s D and nucleotide diversity p. Tajima’s D is used to detect balancing selection, although is also associated with demographic effects. We used ANGSD (v 0.931-6-g18de33d, Korneliussen et al. 2014) to estimate Tajima’s D and nucleotide diversity within 250kb windows across the genome. We considered only reads with a mapping quality above 30 and positions covered by at least one read in 50% of individuals.

We compared the distribution of both metrics for windows detected by each of the three methods to the rest of the genome using a Wilcoxon rank sum test. Because patterns of genetic diversity can covary with recombination rate, we performed the same analysis but this time comparing regions of similar recombination rate. We estimated recombination rates using a linkage map from Rastas et al. (2016). We lift over the coordinates of their SNPs by aligning a 500bp windows around each of their SNPs to our version of the reference genome using bwa-mem (Li 2013). Then, we used Mareymap (v 1.3, Rezvoy et al. 2007) to estimate the recombination rate by linear interpolation across the reference genome (loess interpolation with span = 0.2 and degree = 2). With this, we classified each region of our dataset as associated with low (recombination rate under 1cm/Mb), medium (between 1cM/Mb and 5cM/Mb) or high recombination rates (above 5cM/Mb). This classification was the one used to perform a Wilcoxon rank sum test between the Tajima’s D and p of significant and nonsignificant windows of similar recombination rates.

We identified four genomic regions with putative intra-locus sexual conflicts that are associated with positive Tajima’s D and elevated nucleotide diversity (see Results). To determine if there was an association between significant SNPs in those regions and balancing selection, we estimated Tajima’s D in windows of 25kb (5kb sliding windows) in those regions following the same procedure. Since recent balancing selection can also lead to patterns of increased linkage disequilibrium (LD) (Fijarczyk and Babik 2015), we used plink (v1.90b5.3, Gaunt et al. 2007; Chang et al. 2015; www.cog-genomics.org/plink/1.9/) to estimate LD between all pairs of SNPs within these four genomic regions with putative intra-locus sexual conflicts and their surroundings (we included 250kb flanking regions). To simplify the representation of LD patterns, we used a custom python script to estimate the distribution of pairwise LD for windows of 25kb and represented only the 99^th^ quantile of this distribution. We also used bedtools (v 2.30.0, Quinlan and Hall 2010) to extract genes that were within 1kb or less of a significant SNPs and used the database GeneCards (Stelzer et al. 2016) to search for known functions of these genes. If no functions were available, we used blastX (Altschul et al. 1990) against the uniprot database to identify potential functions by similarity. We then screened each of these genes for potential duplication onto sex chromosomes by calculating F:M coverage bias variation along the gene and within 1kb flanking regions using samtools (Li et al. 2009).

## Supporting information

Supplementary Figures, tables and discussion

## Acknowledgements

The authors are thankful to I. Caza-Allard, E. Reni-Nolin, S. Delaive and F-A. Deschenes-Picard for their help during fieldwork, as well as S. Bernatchez and R. Bouchard for their occasional help. We also thank A-L. Ferchaud and M. Laporte for their advice on analysis, Q. Rougemont for its help in SNP calling and filtering, C. Hernandez and A. Perreault for their help with labwork, and A. Xuereb for proofreading the manuscript. We also thank N. Aubin-Horth for her advice on field and lab work, various exchanges, and in particular for her help around oral communications of this work. Sequencing was performed at the Centre d’Expertise et de Services Genome Québec (Montréal, QC Canada). This research was funded by a Fonds de recherche du Québec - Nature et technologies grant to L. Bernatchez and N. Aubin-Horth. The project is part of the Ressources Aquatiques Québec (RAQ) research program.

## Data availability

The paired end sequencing reads and scripts for analyses and figure reproducibility will be made available upon acceptance.

