## Supplementary Figures, tables and discussion for "Searching for intra-locus sexual conflicts in the three-spined stickleback (*Gasterosteus aculeatus*) genome"

**Supplementary table:**

**Supplementary table 1: summary of sequencing output, filtering steps and SNPs calling in 50 female and 49 male three-spined stickleback.**

**Supplementary table 2: summary of whole genome and chromosome level Redundancy Analysis with sex as explanatory variable.**

Significant p-value indicate that sex explain more variance in the dataset than expected by chance (ChrIX and ChrXII), R2-adj correspond proportion of variance explained by sex.

**Supplementary table 1:**

| | **Number of reads (min-max)** |
|---|---|
| Total number of reads | 37 676 217-105 776 604 |
| % of reads mapped | 0.998793 - 0.999420 |
| Final mean coverage | 12.08128; SD = 3.193991 |
| | **Number of SNPs** |
| Raw number of SNPs | 12 816 669 |
| % of SNPs of bad quality | 36,4% |
| % of SNPs of low coverage | 4,6% |
| % of SNPs with low maf | 45,0% |
| Final number of SNPs | 1 785 441 |

**Supplementary table 2:**

| Scale | P-value | R2-adj |
|---|---|---|
| Whole genome | 0.447 | 1.10E-05 |
| ChrI | 0.938 | -2.54E-04 |
| ChrII | 0.66 | -7.87E-05 |
| ChrIII | 0.121 | 2.50E-04 |
| ChrIV | 0.793 | -1.80E-04 |
| ChrV | 0.194 | 1.90E-04 |
| ChrVI | 0.192 | 1.89E-04 |
| ChrVII | 0.482 | -5.52E-06 |
| ChrVIII | 0.709 | -1.34E-04 |
| ChrIX | 0.046 | 3.61E-04 |
| ChrX | 0.834 | -1.78E-04 |
| ChrXI | 0.887 | -2.49E-04 |
| ChrXII | 0.019 | 3.74E-04 |
| ChrXIII | 0.131 | 2.08E-04 |
| ChrXIV | 0.322 | 8.05E-05 |

| | | |
|---|---|---|
| ChrXV | 0.132 | 2.27E-04 |
| ChrXVI | 0.809 | -1.75E-04 |
| ChrXVII | 0.794 | -1.76E-04 |
| ChrXVIII | 0.231 | 1.48E-04 |
| ChrXX | 0.458 | 1.18E-05 |
| ChrXXI | 0.708 | -1.35E-04 |

**Supplementary Figures:**

**Supp figure 1: Population structure: axes 3 and 4 of a Principal Component Analysis**.

Two outlier individuals with high relatedness are not shown. Ellipses represent the 95% confidence interval of point distribution. Red (circle and ellipse) represent females, blues (triangle and ellipse) represent males

**Supplementary figure 2: Summary of the potential inversion on chrIX**.

A) mds1 score of chrIX from the local pca; the vertical line delimits the putative inversion. B) PCA on the region of the putative inversion. C-D) Variation in heterozygosity and mean depth of sequencing across the inversion for each of the three groups identified on the PCA. E) Intersex Fst across the inversion for all samples (global) and the three groups identified on the PCA.

**Supplemental figure 3: Summary of the potential inversion on chrXVI**.

A) mds2 score of chrXVI from the local PCA; the vertical line delimits the putative inversion. B) PCA on the region of the putative inversion. C-D) Variation in heterozygosity and mean depth of sequencing across the inversion for each of the three groups identified on the PCA. E) Intersex Fst across the inversion for all samples (global) and the three groups identified on the PCA.

**Supplemental figure 4: Summary of the potential duplication on chrXXI.**

A) mds3 score of chrXXI from the local PCA; vertical line delimits the putative duplication. B) PCA on the region of the putative duplication. C-D) Variation in heterozygosity and mean depth of sequencing across the duplication for each of the two groups identified on the PCA. E) Intersex Fst across the duplication for all samples and the two groups identified on the PCA.

**Supp figure 5: Venn diagram of significant SNPs detected with Fisher test, Intersex Fst null model and in the 99% of the Fst distribution.**

We considered a SNP as significant in the Intersex Fst methods if it was identified by all three methods (1478 SNPs).

**Supp figure 6: Cumulative Fst applied at the whole genome and chromosomal scale.**

Each grey dot represents the ratio of the number of SNPs with a significant Fst in our dataset and in a permutation at A) the whole genome scale (black dot represent the mean and its 95%

confidence interval) and B) chromosomal level. At the chromosomal scale, red, green, and blue dots respectively represent the average ratio at the 95, 99 and 99.9 $F_{st}$ quantile with their 95% confidence interval. At both scales, we considered significant enrichment for intra-locus sexual conflict if the mean is >=1 and 1 is not included in the confidence interval.

**Supplementary Figure 7: Association between intra-locus sexual conflict and nucleotidic diversity when controlling for recombination rate.**

Distribution of nucleotidic diversity for each detection method (dashed line) compared to the bulk of the genome (solid line) for each category of recombination rate (low, medium, high, from left to right). From top to bottom: Cumulative $F_{st}$, RDA and SNP-by-SNP. For each method, only category of recombination rate represented by at least 10 windows associated with putative intra-locus sexual conflict are represented; p represents the p-value of a Wilcoxon rank-sum test comparing the distribution of significant and non-significant windows; n represents the number of significant windows.

**Supplementary Figure 8: Intersex coverage bias in genes potentially under balancing selection:**

Coverage bias variation across seven gene associated with intra-locus sexual conflict and potential balancing selection. Region 1Kb around the genes are also represented, as well as exons (grey areas) and significant SNP (red lines).

Supplementary figure 1:

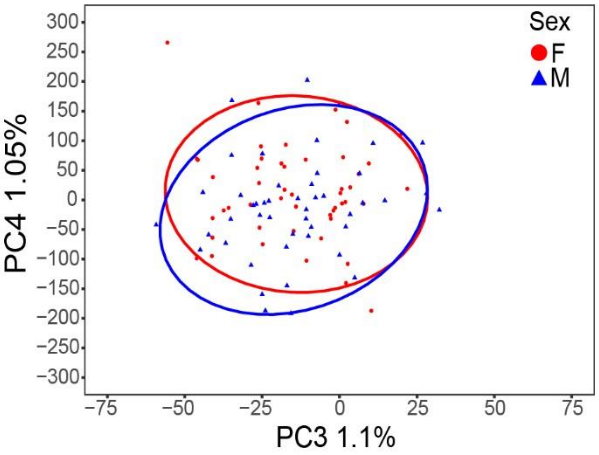


Supplementary figure 2:

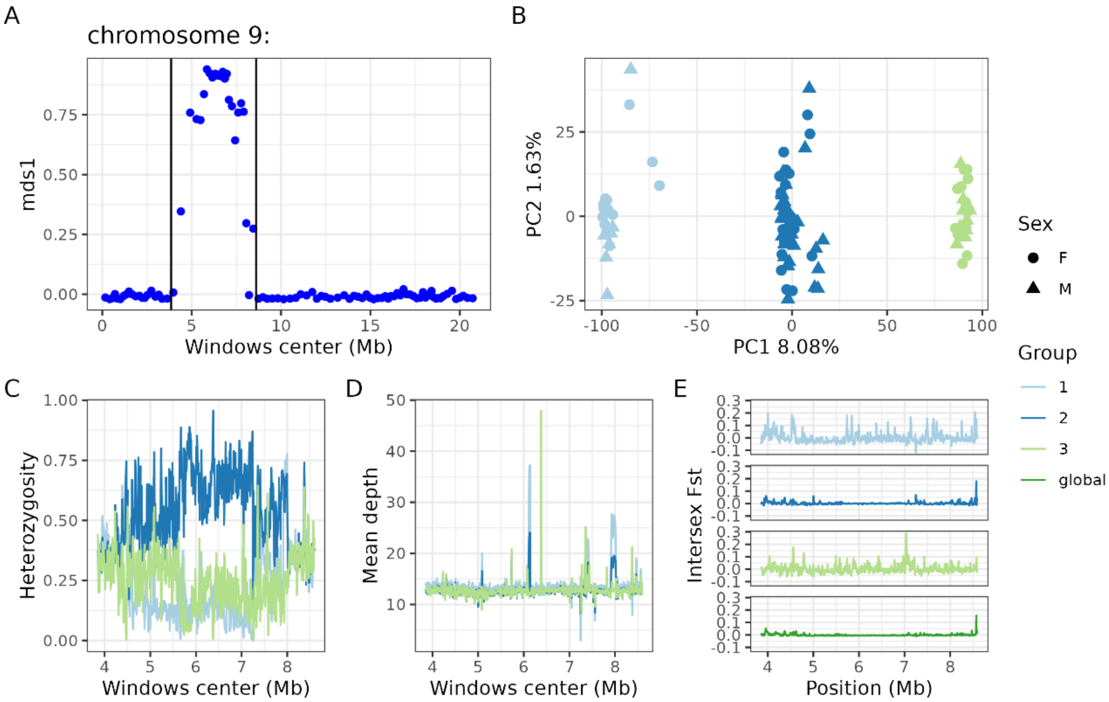

Supplemental figure 3:

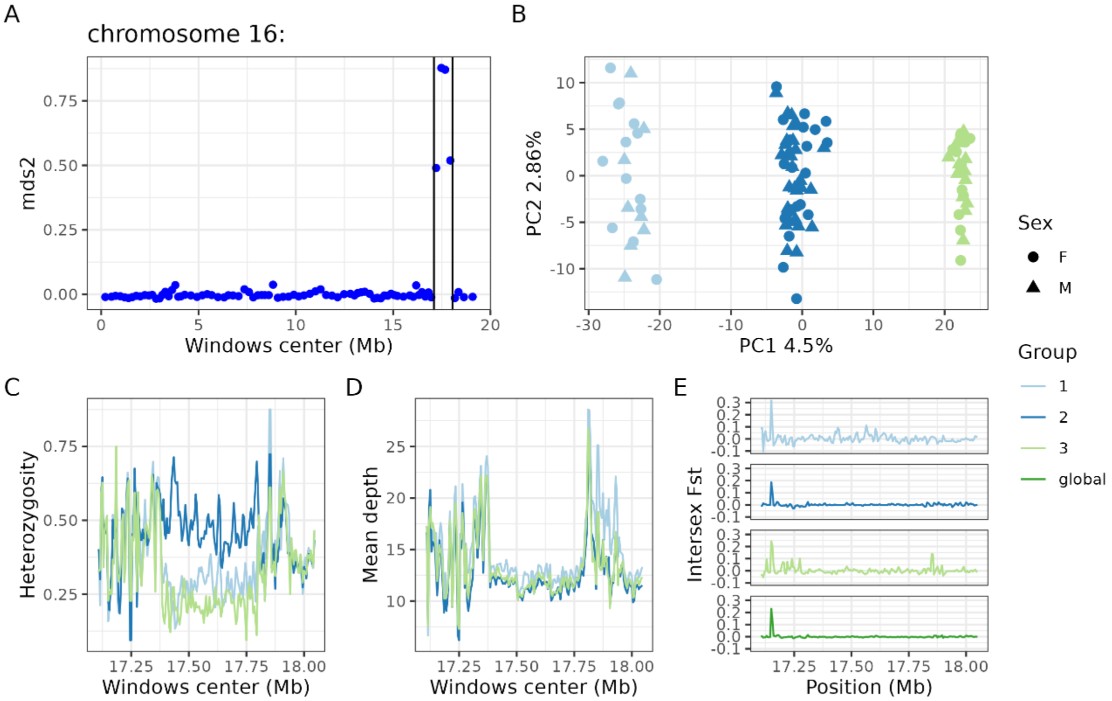


Supplemental figure 4:

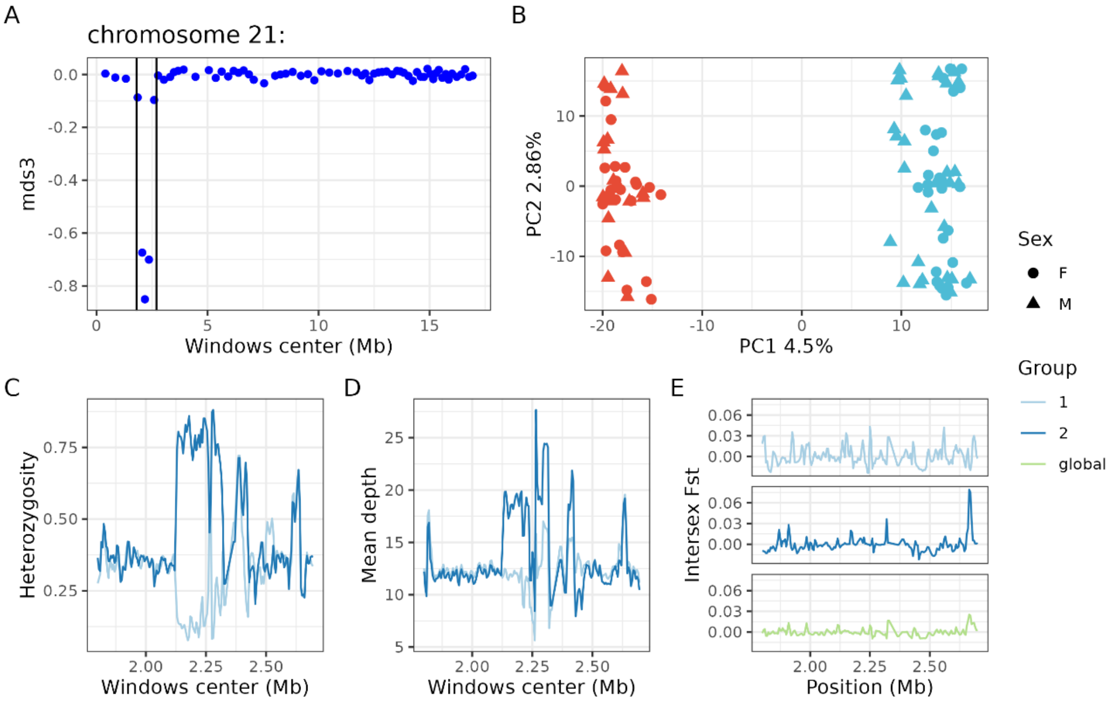

Supplemental figure 5:

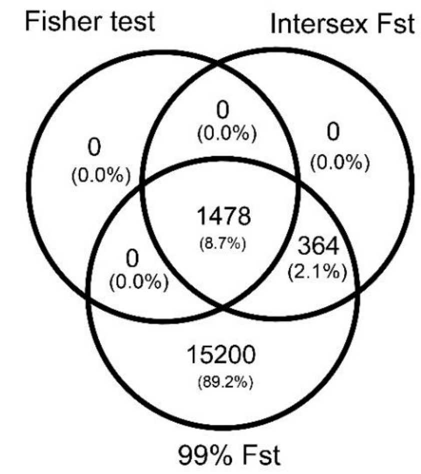


Supplemental figure 6:

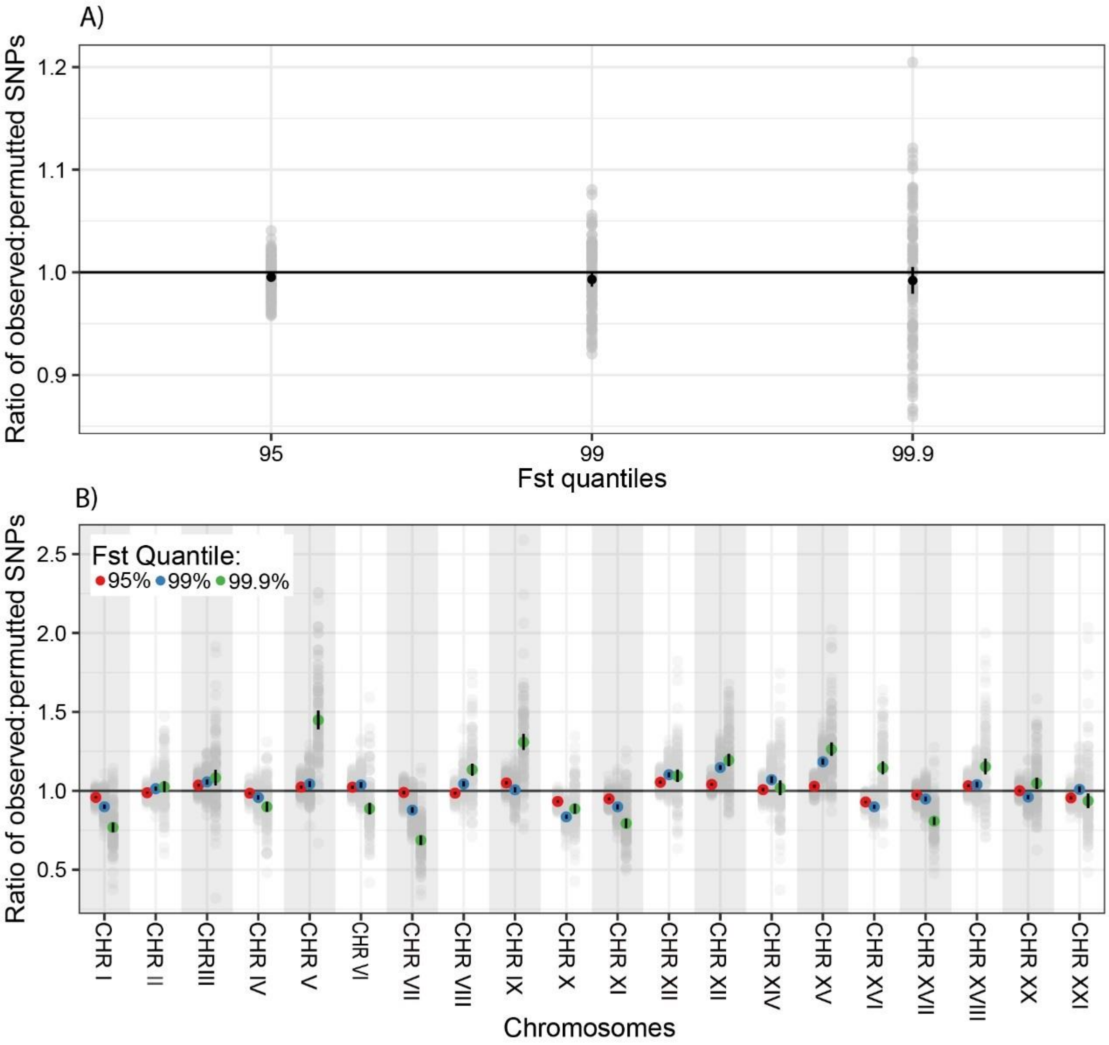

Supplemental Figure 7:

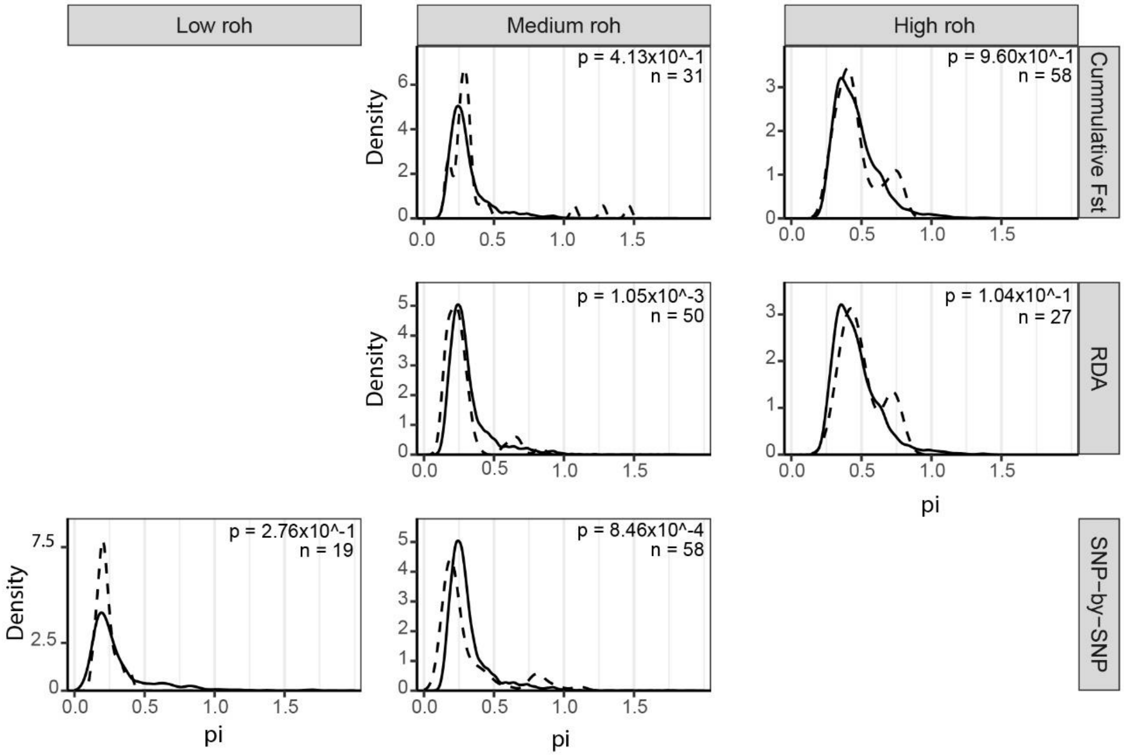

Supplemental Figure 8:

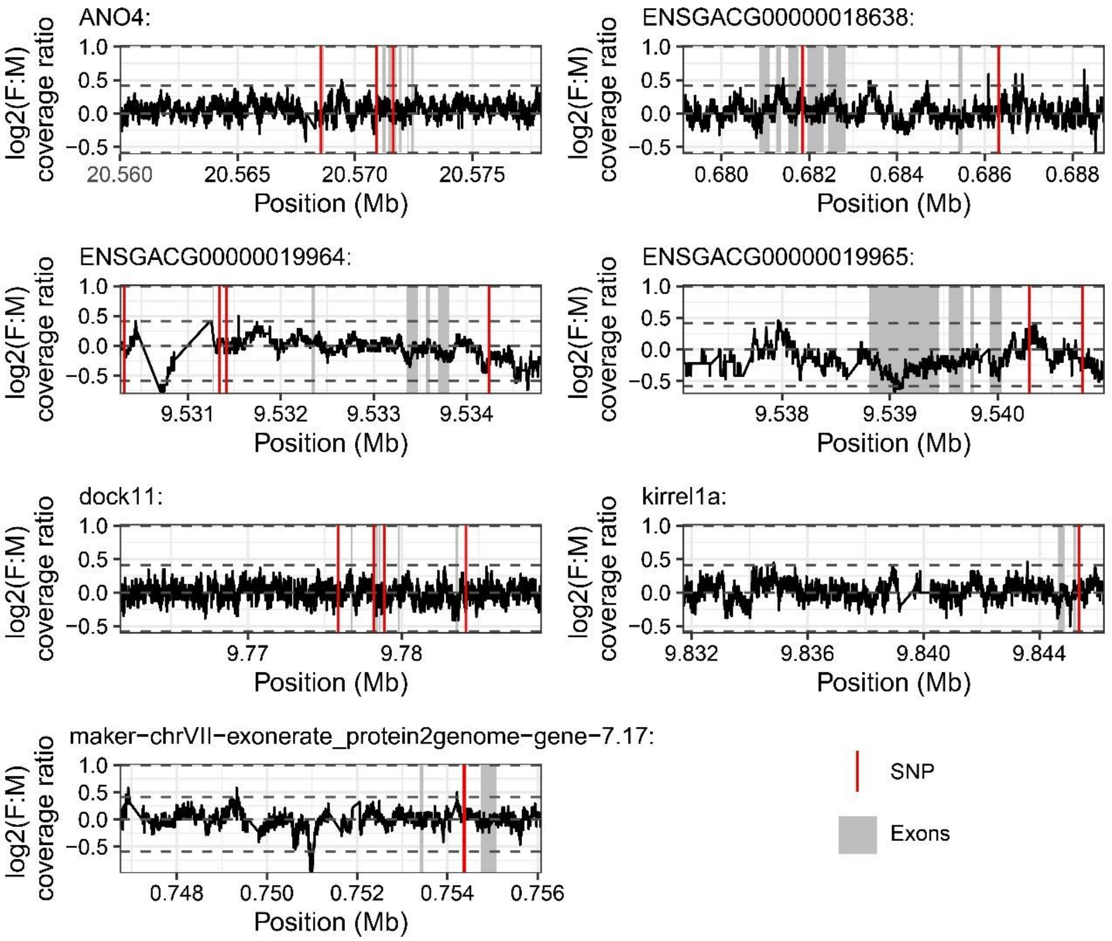


Supplemental discussion: large structural variants in the three-spined stickleback genome.

We identified three potential large structural variants, on chrIX (4.75Mb), chrXVI (0.95Mb) and chrXXI (0.90Mb) (Fig S2, S3, S4). Visual inspection of the NMDS revealed three regions with highly correlated PCA across the genome (Fig S1A, S2A, S3A). To further investigate these regions, we ran another PCA of each region (supp Figure 1B, 2B, 3B). For chrIX and chrXVI, the PCA revealed three distinct groups (kmeans assignment of 98.6% and 99.2% respectively, and PC1 respectively explains 8.08% and 4.5% of the total variance), with the group in intermediate position being more heterozygous than the other groups (Fig S1C and S2C) and no difference in sequencing coverage for the three groups (Fig S1D and S2D). These observations are concordant with a polymorphic inversion forming two haplogroup segregating in the population (group 2 being heterozygous for the two different haplogroup). The PCA performed on chrXXI revealed two groups (kmeans assignment of 98.7%, and PC1 explains 4.5% of total variance, supp figure 3C), with group 2 showing increased heterozygosity and coverage (supp. Figure 3C, D), which is more consistent with a potentially large, duplicated region that could be missing in the reference genome. There was no deviation from Hardy Weinberg equilibrium for both chrIX and chrXVI (pvalue of Chi$^2$ test: chrIX =0.42, chrXVI =0.42), which was not tested for chrXXI since the heterozygous and duplicated groups could not be distinguished. Also, males and females were represented at the same frequency for each of these 3 chromosomes (pvalue for Chi$^2$ test of 0.36, 0.77 and 0.35, respectively). We observed no Fst differentiation between sexes (Fig S1E, S2E, S3E) for these three structural variants, except for a peak of Fst in the third group of the inversion on the chrIX, which is associated with higher sequencing coverage in females (pval) and could reflect the occurrence of a duplication on or from sexual chromosomes. We also observed peaks of Fst for the putative inversion on chrXVI, but because they are located near the breaking points of the inversion, which can be complex to assemble, we did not consider them further. While these structural variations might be of interest for other evolutionary questions, they do not seem to play a role in intra-locus sexual conflict in our study population and were therefore discarded from further analyses.
